# Adaptive synaptic plasticity maintains CRH neuron output during chronic glucocorticoid exposure

**DOI:** 10.1101/2020.04.27.064725

**Authors:** Neilen Rasiah, David Rosenegger, Nuria Daviu, Tamás Füzesi, Jessie Muir, Toni-Lee Sterley, Jaideep S. Bains

**Affiliations:** Hotchkiss Brain Institute Department of Physiology & Pharmacology, Cumming School of Medicine, University of Calgary, 3330 Hospital Drive NW, Calgary, AB, T2N 4N1, Canada; McGill University Department of Psychology 1205 Dr. Penfield Avenue, Montreal, QC H3A 1B1, Canada

## Abstract

An increase in circulating glucocorticoids (CORT) is an essential part of the response to stress. Sustained elevations of CORT, however, have dramatic consequences on behavior, endocrine systems and peripheral organs. Critically, they dampen the endocrine response to acute challenges and decrease intrinsic excitability of corticotropin-releasing hormone neurons in the paraventricular nucleus (CRH^PVN^), suggesting key circuits may be less responsive to stress. Here, we make the surprising discovery that CRH^PVN^ neurons harness a form of adaptive synaptic scaling to escape the persistent negative feedback pressure from CORT and maintain stable output *in vivo*. Specifically, there is an increase in glutamatergic drive to these cells that is mediated by a postsynaptic, multiplicative increase in synaptic strength. These findings suggest that dysfunctions associated with chronic stress may not be due to the primary actions of CORT, but instead reflect the emergence of synaptic adaptations as networks seek to re-establish intrinsic activity setpoints.

## Introduction

In vertebrates, threats to survival initiate defensive responses that include rapid behavioral changes and slow endocrine responses. The endocrine response to stress is controlled by corticotropin-releasing hormone neurons in the paraventricular nucleus of the hypothalamus (CRH^PVN^) at the apex of the hypothalamic adrenal pituitary axis (HPA)^1^. The activation of these neurons is both necessary^1^ and sufficient^2^ to initiate the release of glucocorticoids (CORT) from the adrenal glands. CORT serves multiple functions, including a critical role as the effector of a negative feedback loop that decreases excitability in circuits that drive its release^3,4^.

This exquisite balance between stress, CORT and negative feedback is often threatened. For example, chronic elevations in CORT, termed allostatic load^5^ are evident in disorders like Cushing’s disease^6^, in neuropsychiatric disorders like depression^7,8^ and in patients treated with exogenous glucocorticoids for autoimmune conditions, inflammatory states, and advanced cancers. Although persistent elevations in CORT are prevalent in large clinical cohorts, we understand very little about the consequences for neural stress circuits. This is highlighted by observations that although chronic CORT administration blunts the acute hormone response to stress^9^, it does not alter the reactivity of CRH^PVN^ neurons to acute stress^10^.

In order to understand the consequences of long-term perturbations in circulating CORT on the brain, we need to consider both the actions of CORT itself, but also the reactions of the brain in response to this change in circulating hormone. All biological systems use multiple strategies to defend equilibrium^11^. This homeostatic principle has been studied extensively at the level of large scale physiological processes such as blood pressure, heart rate, temperature and hormone concentrations. But homeostatic principles are also evident in neural networks that utilize multiple strategies to adapt in the face of enduring changes in activity^12–15^. For example, cultured cortical and hippocampal neurons show synaptic adaptations following persistent increases or decreases in activity^12,16,17^. *In vivo,* these synaptic adjustments re-establish activity of visual cortical neurons in the visual cortex following chronic dark exposure^18,19^. Whether physiological systems use similar synaptic adaptations to defend activity setpoints, however, is not well understood.

Here we hypothesized that adaptive synaptic changes in the PVN allow CRH^PVN^ neurons to defend an activity setpoint during persistent CORT negative feedback. By combining a regimen of CORT self-administration with fiber photometry and *ex vivo* electrophysiology, we demonstrate that CORT induced decreases the intrinsic excitability of CRH^PVN^ neurons are balanced by a compensatory upscaling of glutamate synapses allowing this system to remain stable during a chronic challenge.

## Results

### Mechanisms of CORT feedback are maintained after 7 days CORT

To assess how CRH^PVN^ neurons respond to persistent feedback, CRH-CretdTomato mice had their drinking water replaced with a solution containing 25µg/ml CORT dissolved in 0.95% ethanol (EtOH) for 7 days^9,20^. This method applied over several weeks increases circulating CORT during the night phase, and blunts the CORT response to forced swim test (FST)^9,20^, but whether CORT dysregulates this system after only 7 days has not been tested. We first confirmed that the drinking water solution resulted in elevated CORT levels. Tail vein CORT was taken from naïve, EtOH (vehicle), and CORT-treated animals during the dark phase (8pm) on the 7^th^ day (Supplementary Fig. 1a). Circulating CORT was significantly elevated in animals whose drinking water had been replaced with CORT. Also, there was no difference in the volume of solution consumed across groups during the 7 days (Supplementary Fig. 1b). To test whether 7 days of exogenous CORT disrupts HPA output, we measured the CORT response to FST. In these experiments, basal plasma was taken from the tail vein 2hrs (10am) before, and immediately after 15min FST. There was no difference in basal CORT between groups, but animals drinking exogenous CORT for 7 days failed to mount a CORT response to FST (Fig. 1a). This is consistent with blunted HPA activity, and is coherent with previous reports using similar CORT protocols over multiple weeks^9,21^.

**Fig. 1:**
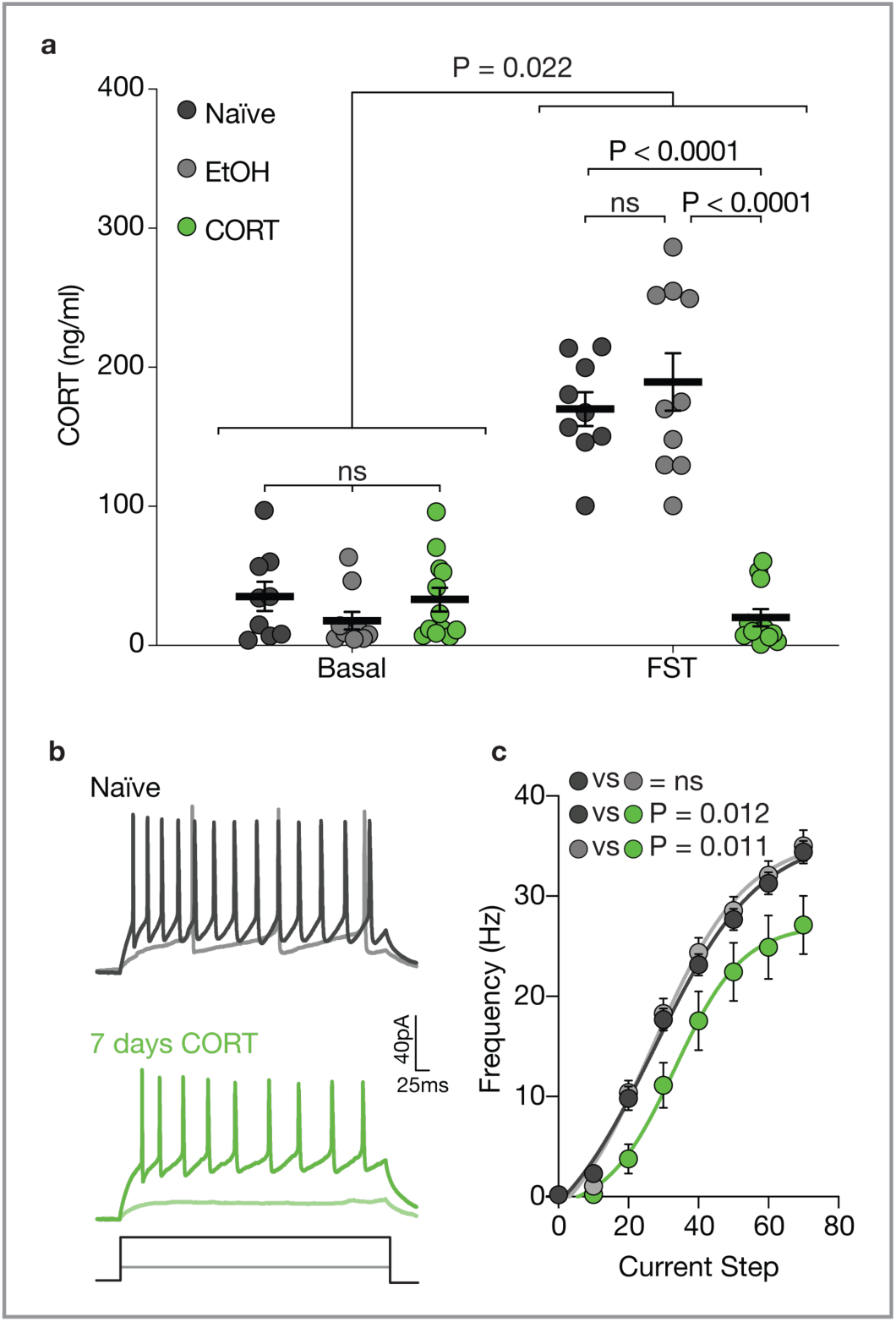
7 days CORT blunts HPA and decreases intrinsic excitability. a) HPA response to stress is blunted following 7 days CORT. A two-way ANOVA with the following factors was used to compare the CORT response to FST: CORT response (F (1, 28) = 120.6, P < 0.0001), treatment (F (2, 28) = 29.05, P < 0.0001), interaction (F (2, 28) = 43.70, P < 0.0001). FST resulted in a significant CORT response in naïve and EtOH treated animals. For naïve (basal, 35.27 ± 10.46ng/ml vs FST, 170.0 ± 12.30ng/ml, N = 9, P < 0.0001) and EtOH-treated (basal, 17.88 ± 6.39ng/ml vs FST, 189.6 ± 20.69ng/ml, N = 10, P < 0.0001), whereas no CORT response was seen in CORT treated animals (basal, 33.03 ± 8.60ng/ml vs FST, 20.08 ± 6.10ng/ml, N = 12, P = 0.75). FST induced a similar rise in CORT among naïve and EtOH animals that was greater than that seen in CORT-treated animals (naïve vs 7 days CORT, P < 0.0001, EtOH vs 7 days CORT, P < 0.0001, and naïve vs EtOH, P = 0.48). There was no difference in basal CORT levels between groups (naïve vs 7 days CORT, P = 0.99, EtOH vs 7 days CORT, P = 0.60, and naïve vs EtOH, P = 0.56). b) Current-clamp recordings from naïve, EtOH, and CORT-treated mice. c) F – I plot showing response to successive current steps. A two-way ANOVA with the following factors was used to compare the F - I plot: Current step (F (2.08, 120) = 590, P < 0.0001), treatment (F (2, 58) = 4.42, P = 0.016), interaction (F (14, 406) = 2.53, P = 0.0018). 7 days CORT resulted in a downshift of the F - I curve when compared to naïve (P = 0.012), and EtOH (P = 0.011). There was no difference between naïve and EtOH (P = 0.95).

CRH^PVN^ neuron activity is sufficient and necessary for CORT production^2^, and CORT feedback after acute stress^3^. Multiple weeks of CORT feedback has been shown to have similar effect^21^ but whether the same effect is seen after 7 days CORT feedback has not been tested. Acute brain slices were prepared to directly assess electrophysiological properties after 7 days CORT. There was a decrease in the excitability of CRH^PVN^ neurons from mice that had access to CORT compared with naïve and EtOH, and no difference between naïve and EtOH (Fig. 1b, c). This confirms that mechanisms of CORT feedback at CRH^PVN^ remain engaged after 7 days CORT.

### Validation of fiber photometry for repeated recordings *in vivo*

Although our *in vitro* recordings demonstrate a decrease in intrinsic excitability of CRH^PVN^ neurons, previous work using *cFOS* as a proxy measure of excitability, *in vivo*, suggest these neurons remain responsive to stress, even during prolonged periods of CORT feedback^10^. To better understand the effect of persistent CORT feedback on CRH^PVN^ neuron activity, we used *in vivo* fiber photometry to assess GCaMP6s activity in live, freely behaving mice. Fiber photometry has emerged as a relatively accessible technique for interrogating neuronal population activity by measuring GCaMP fluorescence^22,23^. We have previously shown that *in vivo* fiber photometry with GCaMP6s works reliably in CRH^PVN^ neurons^24^. Here we first conducted a series of experiments to validate the quantification of GCaMP6s transients as a measure of neural activity, and assessed signal stability for repeated recordings.

Photometry signals in the cortex are highly sensitive to isolfuorane (ISO) anesthesia^25^. We used this approach to compare activity before and during anesthesia, and used peak detection software to quantify calcium transients^26^ (Muir et al., 2018). We obtained baseline recordings from mice in the homecage, then continued recording during transfer to a chamber infused with 4% ISO and return to the homecage (Fig. 2c). We used two different approaches to analyze the data. First, we assessed signal variability by generating all-points histograms from the photometry traces and then comparing histograms from different experimental conditions. Second, we used a peak detection approach to extract information about the frequency and amplitude of discrete GCaMP6s events before and during anesthesia^26^. Peaks were counted if they were greater than a threshold calculated as 2x the median absolute deviation (MAD). For peak detection during baseline and ISO, the detection threshold was set using the MAD from the baseline region of the trace. This was necessary since anesthesia decreases the MAD. During ISO, there was a decrease in the variability of the signal (Fig. 2d-f) and a decrease the frequency of events (Fig. 2g). Based on these observations, we conclude that assessing the variability of the photometry trace as well as event frequency discrete GCaMP6s events provides a reliable readout of changes in the activity of CRH^PVN^ neurons *in vivo*.

**Fig. 2:**
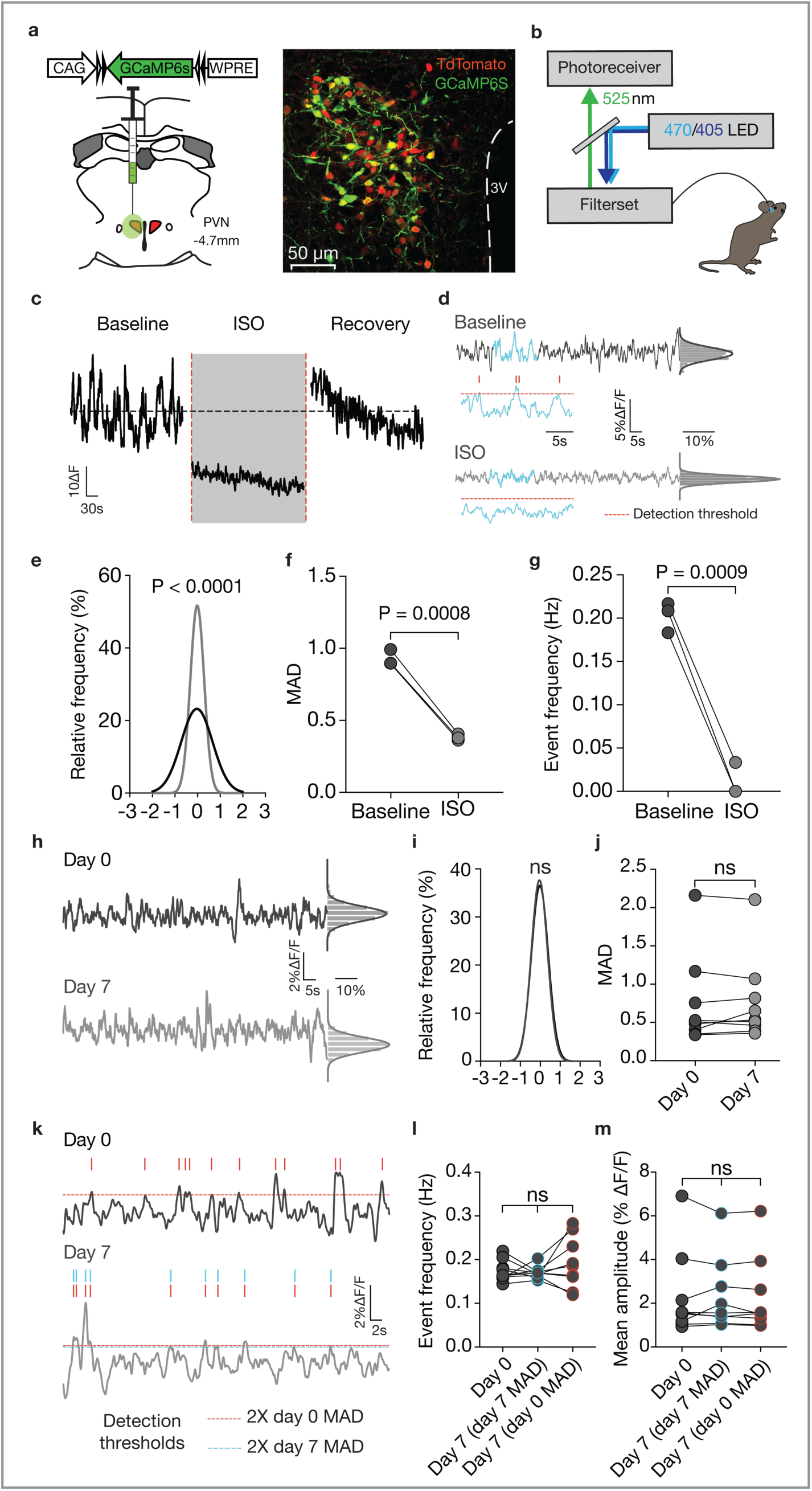
GCaMP6s signal is stable over 7 days. a) Left, showing GCaMP6s viral construct and unilateral injection coordinates. Right, GCaMP6s (green) co-localization with TdTomato-expressing (red) neurons PVN. b) Fiber photometry setup. c) Photometry recordings showing ISO diminishes activity of CRH^PVN^ neurons. d) Peak detection analysis before and during ISO with all-points histogram to the side. Example of detection threshold and peaks are shown below each trace. e) All-points histograms comparing trace variability during baseline and anesthesia (K-S test, P < 0.0001). f) Quantification of MAD before and during ISO (baseline, 0.928 ± 0.0325 vs ISO, 0.382 ± 0.0121, N = 3, paired two-tailed t-test, t(2) = 25.49, P = 0.0015). g) Quantification of the frequency of GCaMP6s events before and during ISO (baseline, 0.203 ± 0.0100Hz vs ISO, 0.0111 ± 0.0111, N = 3, paired two-tailed t-test, t(2) = 23.00, P = 0.0019). h) Recordings from the same animal on day 0 and day 7. i) All-points histogram showing the variability of the trace was unchanged between day 0 and day 7 (K-S test, P = 0.43). j) The MAD was unchanged between day 0 and day 7 (day 0, 0.744 ± 0.197, vs day 7, 0.766 ± 0.183, N = 9, Wilcoxon matched-pairs signed rank test, W = 7.00, P = 0.73). k) Traces highlighting GCaMP6s events. Red-dashed line indicates peak detection threshold determined using day 0 MAD, and the blue-dashed line denotes the detection threshold calculated with day 7 MAD. Ticks above traces signify detected events. l) There was no significant difference in event frequency between day 0 and day 7 using either day 0 or day 7 threshold (day 0, 0.175 ± 0.00799Hz, vs day 7 (with day 7 MAD), 0.170 ± 0.00469Hz, vs day 7 (with day 0 MAD), 0.192 ± 0.0195Hz, N = 9, one-way ANOVA, Tukey correction, F(1.34, 10.7) = 1.02, P = 0.36). m) There was no difference in event amplitude (day 0, 2.32 ± 0.652%∆F/F, vs day 7 (with day 7 MAD), 2.34 ± 0.556%∆F/F, vs day 7 (with day 0 MAD), 2.30 ± 0.580%∆F/F, N = 9, one-way ANOVA, Tukey correction, F(2, 16) = 0.0588, P = 0.94).

Next, we validated the stability of the photometry signal over the course of 7 days by obtaining recordings 7 days apart from the same animal in its homecage (Fig. 2h). There was no change in the all-points histogram (Fig. 2i), or the MAD between day 0 and day 7 (Fig. 2j). This indicates that the signal remains stable, and is suitable for repeated recordings. We then asked whether the peak detection would remain consistent over 7 days (Fig. 2k). While doing this, we considered the possibility that CORT could alter baseline variability, which in turn may confound peak detection parameters. Therefore, it was critical to assess peak frequency and amplitude in an unbiased fashion. Since the peak detection threshold is determined by the MAD, and there is no difference in MAD between day 0 and day 7, we tested whether the peak detection threshold from day 0 could be applied to the day 7 recording without significantly changing the frequency and amplitude of detected events. After 7 days, there was no significant difference in peak frequency (Fig. 2l) or amplitude (Fig. 2m), and applying the day 0 detection threshold to the day 7 recording did not change frequency or amplitude. Altogether, these experiments demonstrate the stability of the GCaMP6s signal over 7 days and indicate this approach is reliable for assessing changes in neuronal activity during repeated recordings.

### CRH^PVN^ neurons remain active during persistent CORT feedback

Photometry recordings were obtained before (day 0) and after the 7^th^ day of CORT feedback in the homecage (Fig. 3a). The variability of the photometry signal was unchanged by CORT (Fig. 3b) and there was no difference in the MAD (Fig. 3c). We assessed the frequency and amplitude of spontaneous events (Fig. 3d) and failed to detect any changes in either frequency (Fig. 3e) or amplitude (Fig. 3f) of GCaMP6s events before and after CORT. These data indicate that despite decreased intrinsic excitability, the overall activity of CRH^PVN^ neurons is not changed by chronic CORT.

**Fig. 3:**
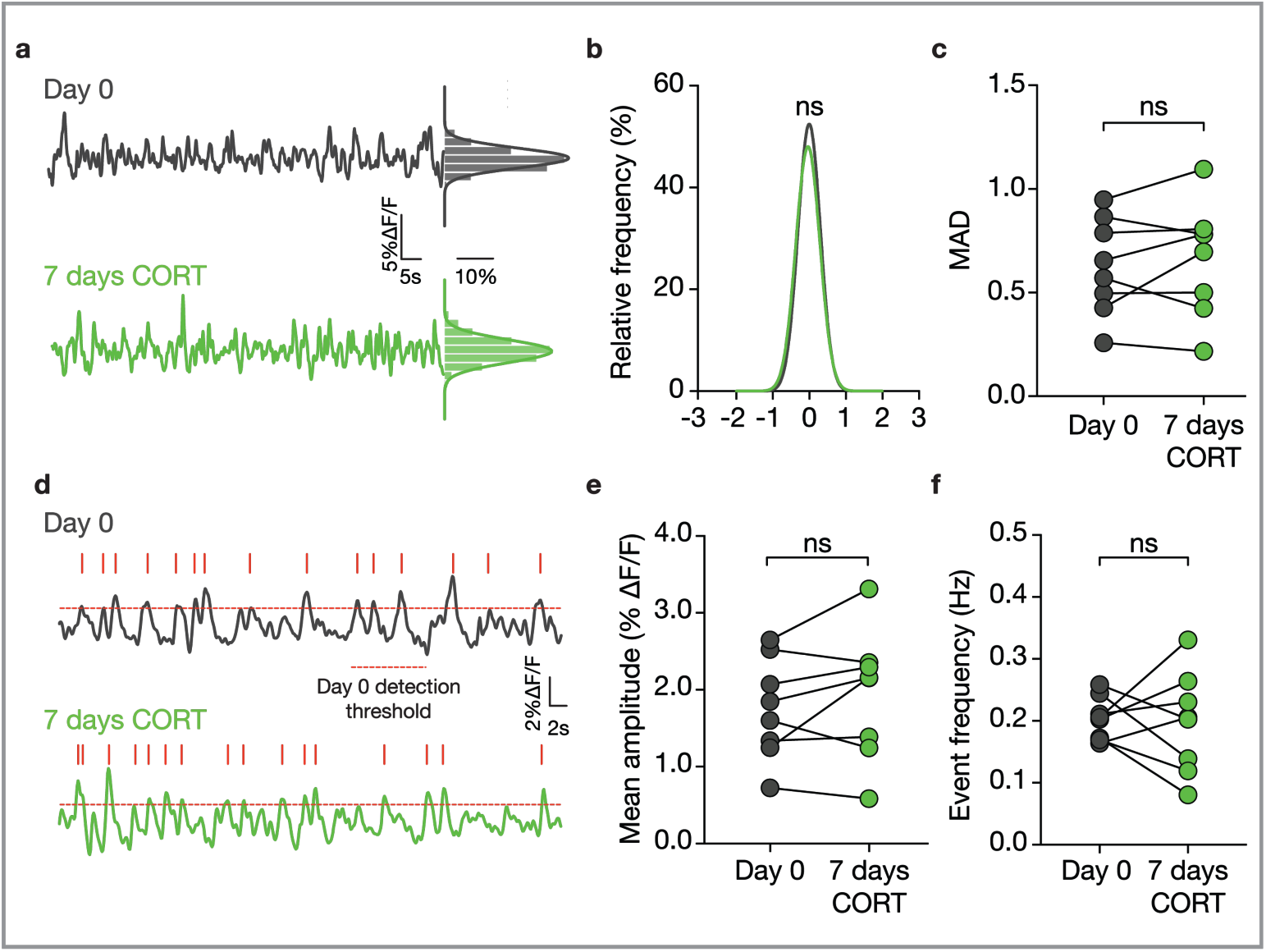
CRH^PVN^ neurons remain active during persistent CORT feedback. a) Traces from the same animal on day 0 and after 7 days CORT with all-points histograms to the side. b) The variability of the trace was unaffected by 7 days CORT as indicated by the all-points histograms (K-S test, P = 0.084). c) The MAD was unaffected by 7 days CORT (day 0, 0.626 ± 0.0828, vs day 7, 0.663 ± 0.0963, N = 8, paired two-tailed t-test, t(7) = 0.761, P = 0.47). d) Traces highlighting GCaMP6s events with red ticks signifying spontaneous events. Red-dashed line indicates peak-detection threshold from day 0. e) 7 days CORT had no effect on peak amplitude (day 0, 1.75 ± 0.23%∆F/F, vs day 7, 1.9 ± 0.28%∆F/F, N = 8, paired two-tailed t-test, t(7) = 1.21, P = 0.27). f) 7 days CORT had no effect on frequency (day 0, 0.204 ± 0.0123Hz, vs day 7, 0.197 ± 0.0288Hz, N = 8, P = 0.82, paired two-tailed t-test, t(7) = 0.242, P = 0.82).

### Global and multiplicative increase in glutamate synaptic strength following 7 days CORT

We hypothesized that persistent CORT feedback results in compensatory synaptic changes that override dampened excitability. To interrogate the effect of 7 days CORT on synaptic drive, we recorded α-amino-3-hydroxy-5-methyl-4 isoxazolepropionic acid receptor (AMPAR) currents with whole-cell electrophysiology. We found that 7 days CORT increased the amplitude of miniature excitatory postsynaptic currents (mEPSCs) (Fig. 4a, b), and no effect on mEPSC frequency (Fig. 4c). Next, we assessed spontaneous inhibitory postsynaptic currents (sIPSCs) (Supplementary Fig. 2a). There was no change in the amplitude or frequency of sIPSCs from γ-aminobutyric acid (GABA) synapses (Supplementary Fig. 2b, c).

**Fig. 4:**
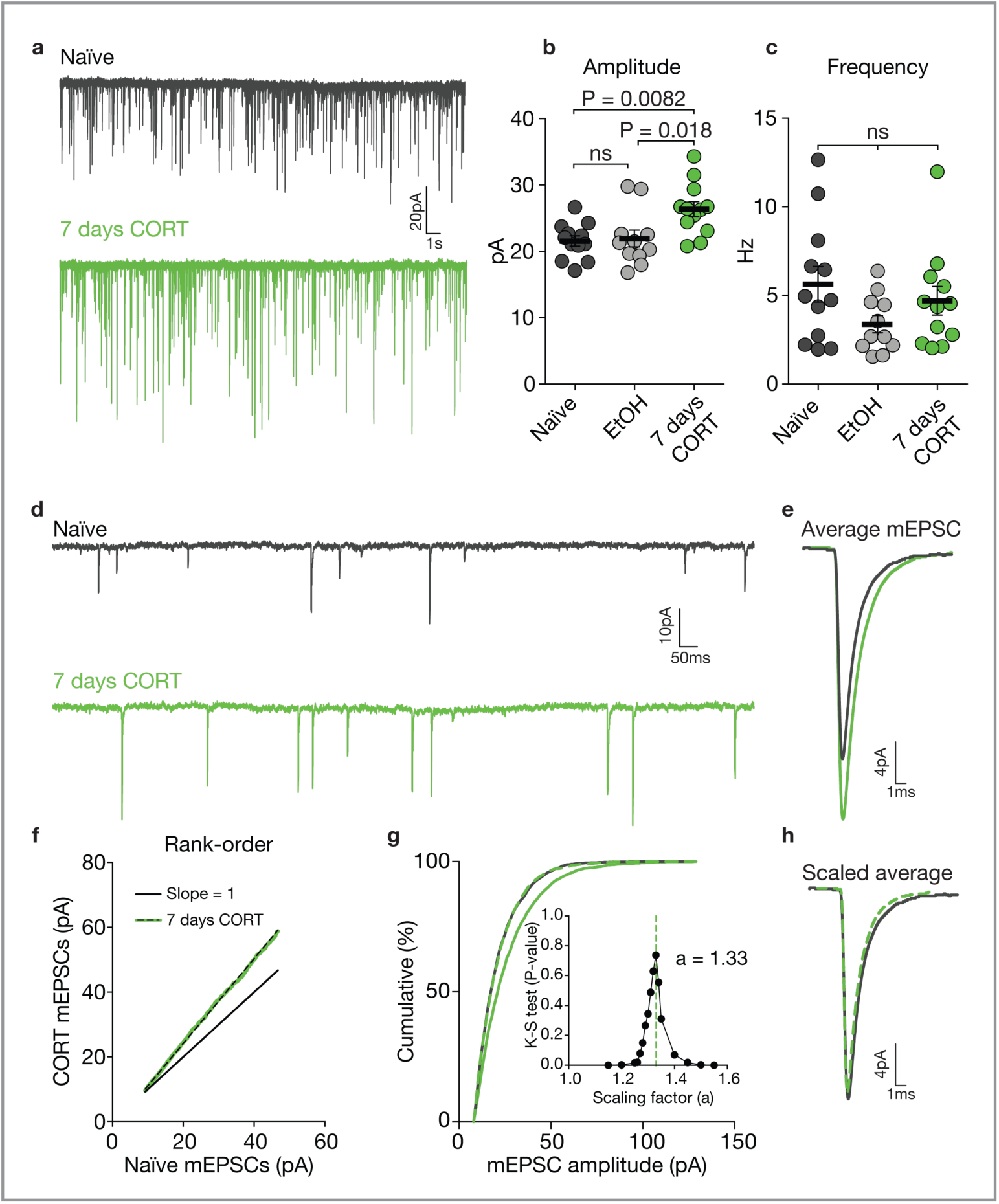
Enhanced excitatory transmission after 7 days CORT. a) Voltage-clamp recordings of mEPSCs from CRH^PVN^ neurons. b) There was a significant increase in mean mEPSC amplitude after 7 days CORT compared to naïve and EtOH (naïve 21.6 ± 0.781pA, n = 12, vs, EtOH, 21.9 ± 1.26pA, n = 11, N = 4, vs 7 days CORT, 26.4 ± 1.14pA, n = 12, one-way ANOVA, Tukey correction, F (2, 32) = 6.34, P = 0.0048). c) 7 days CORT had no effect on mEPSC frequency (naïve, 5.63 ± 1.00Hz, n = 12, vs EtOH, 3.38 ± 0.486Hz, n = 11 vs 7 days CORT, 4.69 ± 0.806Hz, n = 12, one-way ANOVA, Tukey correction, F (2, 32) = 1.91, P = 0.16). d) Traces showing mEPSCs from naïve and 7 days CORT treated mice. e) Average mEPSC from each condition. f) Rank-order plot of mEPSC amplitudes from naïve animals (slope = 1) vs 7 days CORT treated animals (linear regression, Y=1.31X – 2.01, R^2^ = 0.99, P < 0.0001). g) Cumulative distribution showing mEPSC amplitudes from CORT treated animals. The K-S test was used to determine 1.33 as the scaling factor as it yields highest P-value (0.74) (inset). h) Average mEPSC from naïve overlaid with scaled mEPSC after 7 days CORT.

We further characterized the effect of 7 days CORT on excitatory synapses to elucidate a process governing the increase in mEPSC amplitude (Fig. 4d). Averaged mEPSCs from naïve and CORT treated animals are shown (Fig. 4e). A rank-order plot containing mEPSCs recorded from naïve and CORT treated animals was made (Fig. 4f). The linear regression yielded a line with slope = 1.31, and values tightly followed the regression (R^2^ = 0.99). This indicates that the entire distribution of mEPSCs is increased by a factor common to all synapses. The factor was calculated using a validated approach^27^ (Kim et al., 2012), and was determined to be 1.33 (Fig. 4g). The average mEPSC from naïve is shown compared to the average CORT mEPSC scaled down by 1.33 (Fig. 4h). This analysis indicates that 7 days of CORT feedback results in a multiplicative increase in the amplitude of AMPAR synapses that occurs across the entire population. We hypothesize this is an adaptive process that maintains activity of CRH^PVN^ during prolonged CORT feedback.

### Increased postsynaptic AMPARs is likely a mechanism for increased amplitude

Increased mEPSC amplitude may occur through a pre- or postsynaptic mechanism. We first tested for presynaptic mechanisms by assessing vesicular release probability. Whole-cell recordings were taken from CRH^PVN^ neurons while a stimulating electrode was placed in the adjacent neuropil along the paraventricular aspect of the PVN^28^. Two electrically evoked EPSCs (50ms apart) were recorded in the presence of picrotoxin (100µM), and the paired-pulse ratio (PPR) was compared. There was no difference in PPR from naïve and CORT animals (Fig. 5a, b), which indicates that presynaptic release does not contribute to increased mEPSC amplitude. Another potential presynaptic origin for increased mEPSC amplitude could be enhanced quantal packaging of glutamate. Augmented quantal packaging would increase glutamate concentration in the synaptic cleft, and can be tested by comparing the effect of partial AMPAR antagonism has on mEPSC amplitude^29^. The AMPAR antagonist, γ-D-glutamaglycine (γ-DGG) has rapid dissociation kinetics and binds with low affinity^29^. As cleft concentrations of glutamate rise, glutamate should out-compete γ-DGG at the receptor binding site, and γ-DGG will be less effective at antagonizing AMPAR currents^29^. Therefore, if CORT increased quantal packaging, γ-DGG would be less effective at antagonizing AMPAR currents in CORT treated animals. In these experiments, mEPSCs were first recorded in the absence of γ-DGG, then γ-DGG (500µM) was superfused into the recording chamber, and another set of mEPSCs was recorded. γ-DGG has been shown to preferentially block smaller events^29,30^. This can be seen in the normalized cumulative graphs (Fig. 5c). A ratio of mean amplitude before and after γ-DGG shows no significant difference in the effectiveness of γ-DGG antagonism on AMPAR currents between CORT and naïve animals (Fig. 5d). These experiments demonstrate that presynaptic mechanisms do not contribute to increased mEPSC amplitude.

**Fig. 5:**
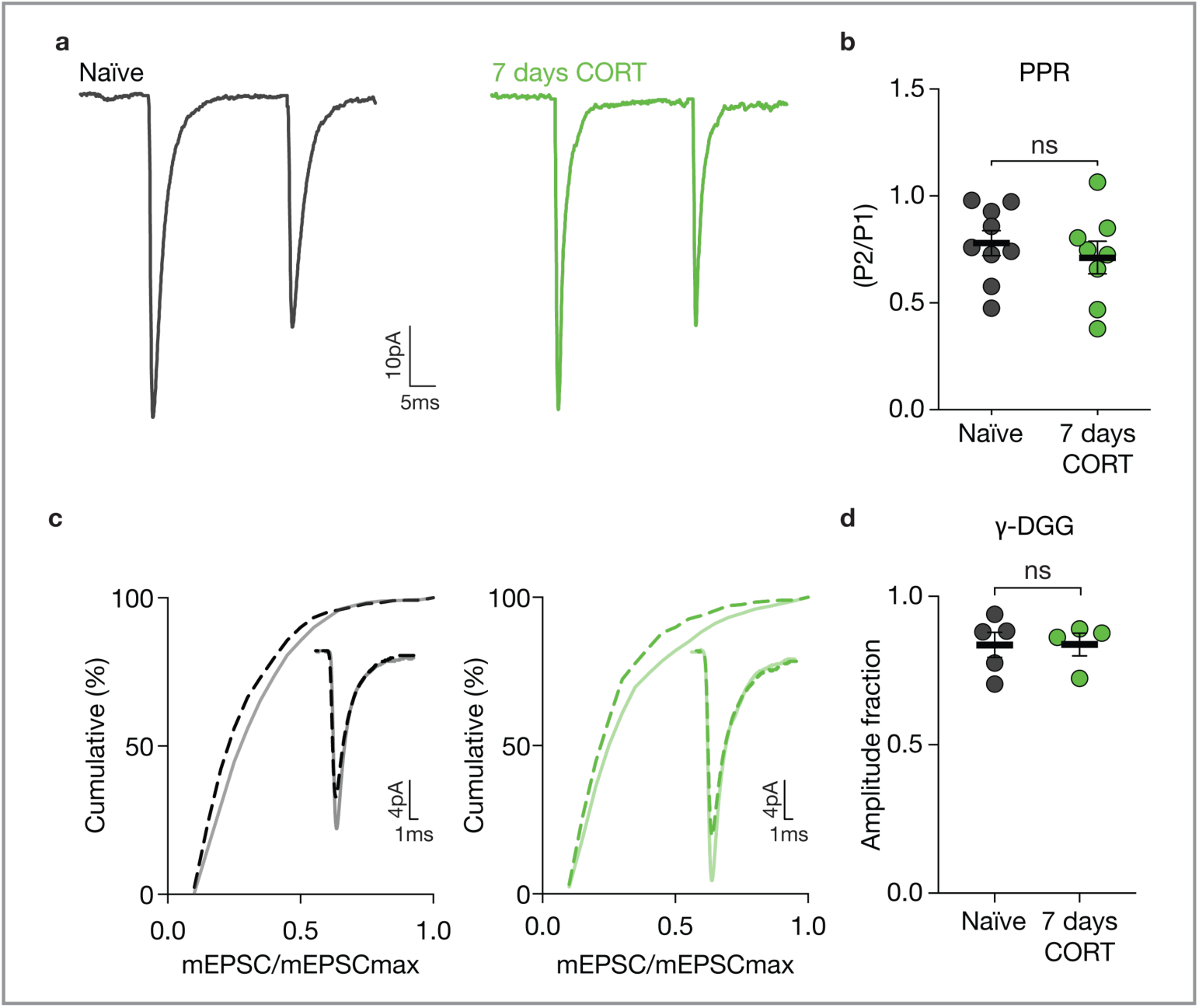
Presynaptic processes do not contribute to enhanced mEPSC amplitude. a) Representative electrically evoked EPSCs. b) There was no difference in PPR after 7 days CORT (naïve, 0.780 ± 0.0582, n = 9, vs 7 days CORT, 0.712 ± 0.0763, n = 8, unpaired two-tailed t-test, t(15) = 0.714, P = 0.49). c) Cumulative histograms of normalized mEPSC amplitudes before and after γ-DGG in naïve (left) and CORT (right) animals. d) There was no difference in the amplitude fraction of mEPSCs from naïve and CORT-treated animals (naïve, 0.839 ± 0.0431, n = 5, vs 7 days CORT, 0.84 ± 0.0391, n = 4, two-tailed Mann-Whitney test, U = 9, P = 0.90).

We next probed for postsynaptic changes that could account for increased mEPSC amplitude following CORT. There was no change in membrane resistance or capacitance after CORT, which indicates that changes in membrane conductance did not alter the fidelity of glutamate currents as they travelled toward the recording electrode after CORT. Also, the access resistance of the patch pipette was not different during recordings of naïve and CORT treated animals (Supplementary Fig 3a). Another possibility is that 7 days CORT enhances conductance of individual AMPAR channels. However, the 10-90 rise time of AMPAR currents between naïve and CORT treated animals was not different (Supplementary Fig. 3b). This indicates that CORT does not change AMPAR conductance. Since there were no changes in intrinsic membrane properties of CRH^PVN^ neurons, mEPSC kinetics, presynaptic release probability, or increases glutamate concentration in the synaptic cleft, we conclude that increased mEPSC amplitude is likely due to the postsynaptic insertion of AMPARs.

### Co-activation of TrkB prevents increased mEPSC amplitude

CRH^PVN^ neurons express tyrosine receptor kinase B (TrkB), the receptor for brain-derived neurotrophic factor (BDNF)^31^. Stress, through GR activation reduces BDNF expression^32,33^ and TrkB activation^31^. BDNF prevents homeostatic synaptic up scaling during activity blockade^17^. TrkB increases the expression of activity-regulated cytoskeletal protein (Arc/arg3.1)^34–36^, which regulates surface AMPARs through endocytic machinery^37^. Arc/arg3.1 overexpression prevents the multiplicative increases in AMPAR amplitude induced by activity blockade, and Arc/arg3.1 knockout in the absence of activity blockade results in a multiplicative increase in synaptic strength^38^. Based on these observations, we hypothesized that increasing TrkB signaling should decrease synaptic scaling at glutamate synapses.

We used a pharmacological approach to increase TrkB signaling during CORT administration. 7,8-dihydroxyflavone (DHF) is a plant derived TrkB agonist that has favorable gut absorption when delivered orally, and can penetrate the blood-brain barrier^39^. One group of animals was given access to a solution containing DHF (160μg/ml) and CORT, and a second group was given the CORT solution as described above. After 7 days of free access to either solution, animals were sacrificed and acute brain slices were prepared for whole-cell recordings (Fig. 6a). The amplitude of mEPSCs in CRH^PVN^ neurons from CORT-treated animals was larger than the amplitude from animals treated with CORT + DHF (Fig. 6a, b). There was no change in mEPSC frequency (Supplementary Fig. 4a). Inclusion of DHF did not change input resistance or capacitance, and the access resistance during recordings was similar between the groups (Supplementary Fig. 4b). The presence of DHF had no effect intrinsic excitability (Fig. 6c, d).

**Fig. 6:**
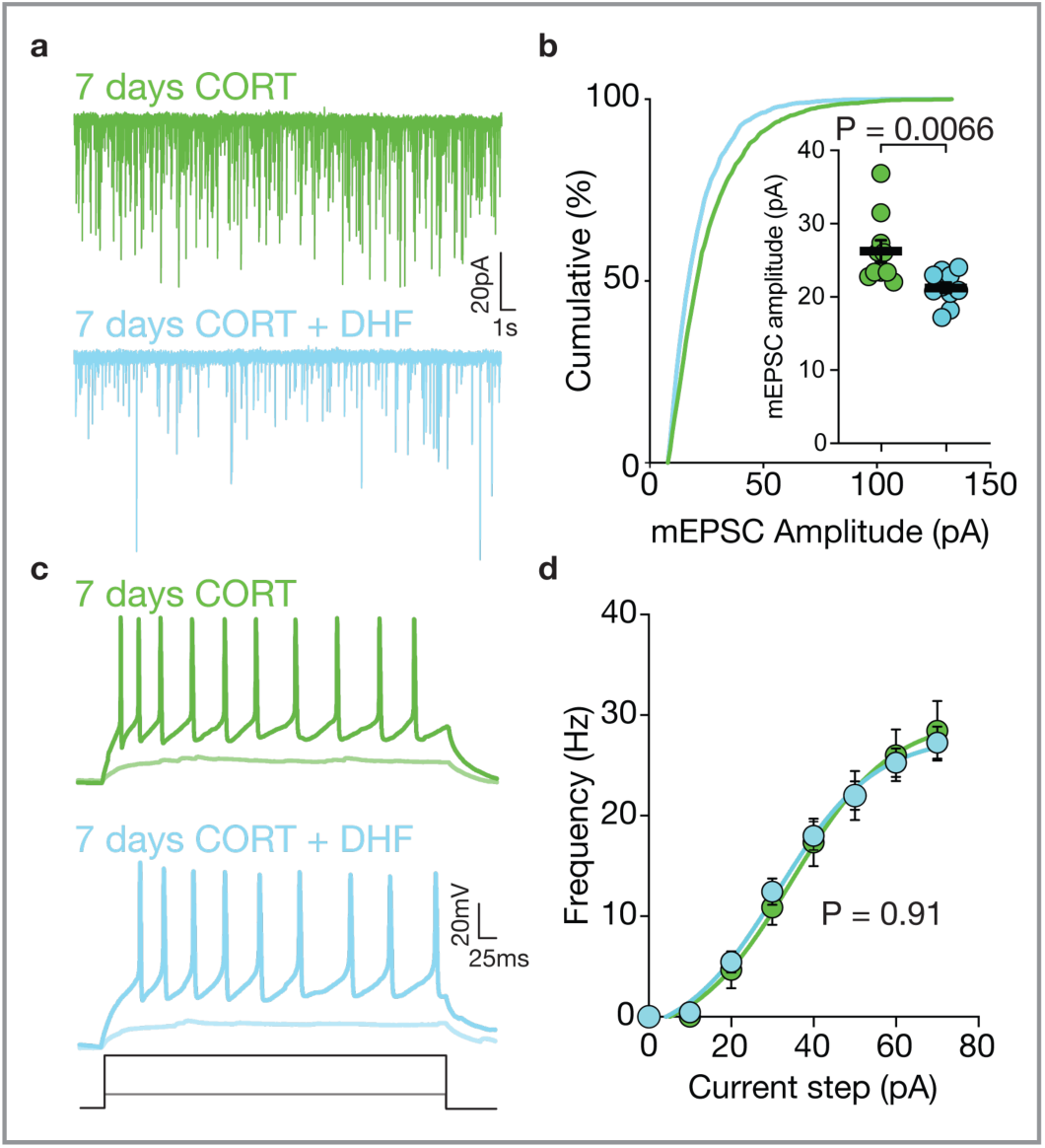
Synaptic scaling blocked by TrkB activation. a) mEPSCs from animals given CORT or CORT + DHF. b) DHF + CORT solution disrupted the multiplicative shift in the mEPSC amplitude distribution (K-S test, P < 0.0001), and the increase in mean mEPSC amplitude (inset) (7 days CORT, 26.3 ± 1.48pA, n = 10, 7 days CORT + DHF, 21.3 ± 0.717pA, n = 10, unpaired two-tailed t-test, t(18) = 3.07, P = 0.0066). A two-way ANOVA with the following factors was used to compare the F-I plot: Current step (F (2.00, 57.84) = 212.00, P < 0.0001), treatment (F (1, 29) = 0.012, P = 0.91), interaction (F (7, 203) = 0.32, P = 0.94). There was no difference between 7 days CORT and 7 days CORT + DHF.

### Adaptive synaptic plasticity maintains CRH^PVN^ neuron activity

We used DHF to probe for the contributions of increased synaptic drive to CRH^PVN^ activity *in vivo*. We repeated photometry experiments before and after 7 days of CORT + DHF while animals were in their homecage (Fig. 7a). Following 7 days CORT + DHF there was a decrease in signal variability, as the all-points histogram showed a kurtosis relative to day 0 (Fig. 7b), and there was a significant decrease in the MAD (Fig. 7c). There was no change in the amplitude of spontaneous GCaMP6s events (Fig. 7d, e), but we did observed a decrease frequency (Fig. 7f). These data indicate that reduced intrinsic excitability in the absence of a concomitant increase in mEPSC amplitude across the entire population of excitatory synapses decreases the output of CRH^PVN^ neurons. This suggests that increased drive at glutamate synapses is an adaptive process that maintains CRH^PVN^ activity during prolonged CORT feedback.

**Fig. 7:**
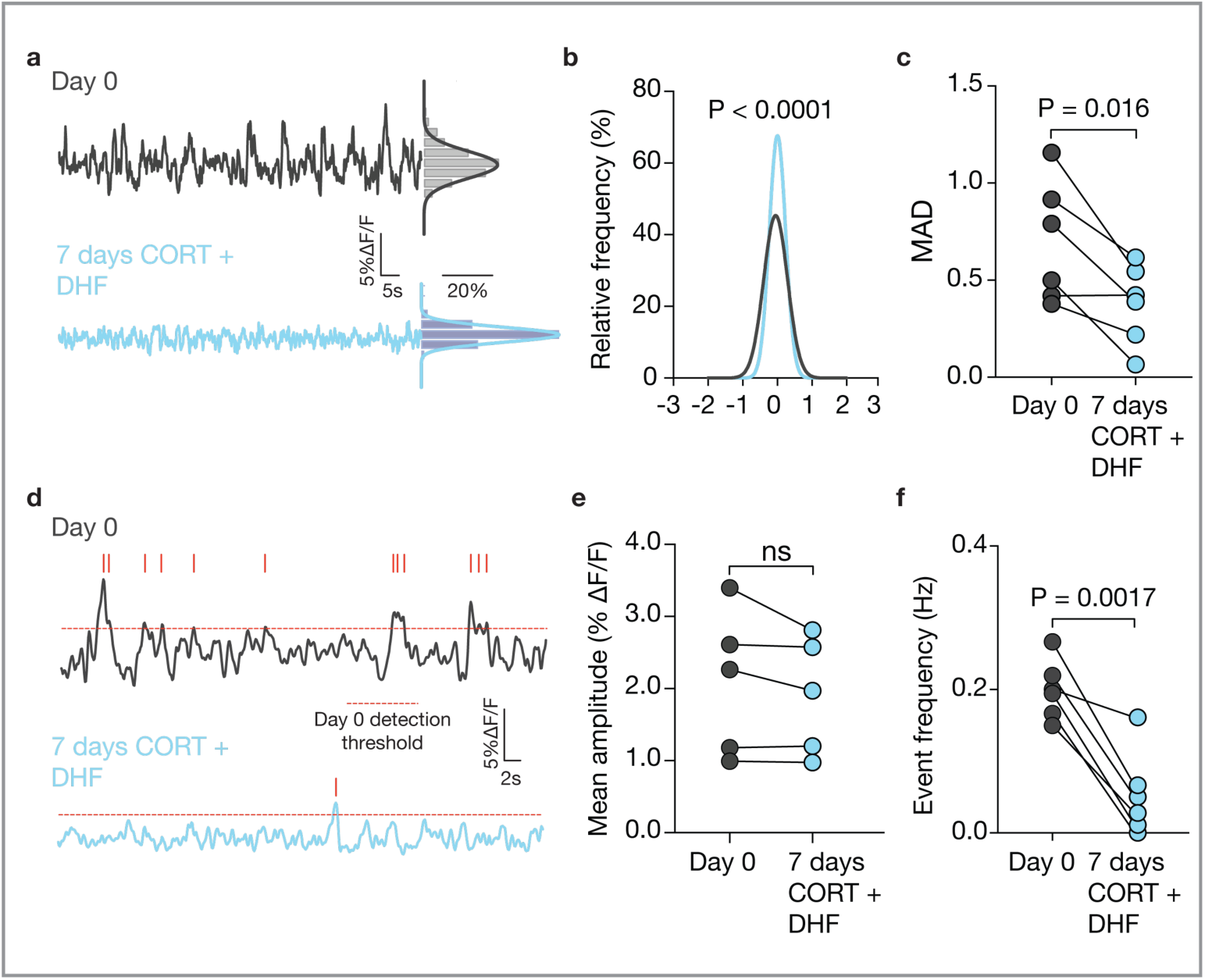
Scaling maintains activity during persistent CORT feedback. a) Baseline traces highlighting decrease in signal variability on day 0 and after 7 days CORT + DHF. b) Consistent with the decrease in variability, there was a kurtosis in the all-points histograms (P < 0.0001, K-S test). c) There was also a significant decrease in the MAD after 7 days CORT + DHF (day 0, 0.694 ± 0.127 vs day 7, 0.376 ± 0.0836, paired two-tailed t-test, t(5) = 3.582, P = 0.016). d) Traces showing GCaMP6s transients with detected peaks. e) There was no difference in the amplitude of GCaMP6s events (day 0, 2.09 ± 0.450%∆F/F vs day 7, 1.91 ± 0.363%∆F/F, N = 5, paired two-tailed t-test, t(4) = 1.577, P = 0.19). One animal was excluded from statistical analysis as there was no detectable events after 7 days CORT + DHF. f) There was a significant decrease in event frequency after 7 days CORT + DHF (day 0, 0.200 ± 0.0168Hz vs day 7, 0.0528 ± 0.0239Hz, N = 6, paired two-tailed t-test, t(5) = 6.11, P = 0.0017).

## Discussion

Here, we show that long-lasting CORT feedback to CRH^PVN^ neurons recruits an adaptive program to multiplicatively increase the strength of glutamate synapses. *In vivo*, the net effect of these changes is to stabilize CRH^PVN^ neuron activity. This is the first demonstration that neural networks responsive to hormonal feedback leverage adaptive mechanisms to overcome feedback. Previous studies show acute CORT feedback decreases activity *in vivo*^4^, yet *cFOS* studies suggest that activity is maintained after chronic feedback^10^. Our results provide stronger evidence that activity is maintained *in vivo*, and provide a synaptic mechanism that allows these neurons to overcome feedback.

Our experimental design, to administer CORT through the drinking water, maintains the rhythmicity of daily circadian CORT release^9^, as CORT levels are low in the morning, but peak during the night (when the animal is active and drinking the solution). After 7 days, this led to a decrease in CRH^PVN^ neuron excitability, which is in line with previous findings that CORT alters activation kinetics of hyperpolarizing K+ channels in CRH^PVN^ neurons following acute stress^3^. While, we do not propose this necessarily as a model of chronic stress, it does allow us to examine the consequences of a single variable within the cascade of changes that occur during chronic stress.

The decrease in intrinsic activity of CRH^PVN^ *in vitro* following 7 days CORT was countered with a multiplicative increase in excitatory synaptic strength. This is reminiscent of the original descriptions of multiplicative synaptic scaling using cultured neurons^12,16,17,40^. A validated approach^27^ was used to determine that the increase in mEPSC amplitude is consistent with a multiplicative synaptic scaling of glutamate synapses. We then probed how this increase in synaptic strength occurs. In our experiments, we were unable to find presynaptic changes such as increased release probability or increased quantal content after 7 days CORT. Therefore, the most parsimonious explanation for our observations is an increase in the number of postsynaptic AMPARs.

We then show that activation of TrkB during CORT feedback prevents the increase in synaptic amplitude. Importantly, the intrinsic excitability of CRH^PVN^ neurons treated with CORT was not affected by the addition of DHF. When assessing CRH^PVN^ neurons *in vivo*, CORT + DHF resulted in reduction of GCaMP6s activity *in vivo*, indicating decreased activity. However, in animals given CORT alone, there was no difference in the signal before and after CORT, indicating that adjustment in synaptic amplitude maintains basal activity in this system. The precise mechanisms through which this occurs remains unresolved, but a likely possibility is that the activation of TrkB receptors induces Arc/arg3.1, which increases the rate of AMPAR endocytosis via interaction with dynamin and endothelin^37^. Alternatively, we cannot rule out an interaction between BDNF and tumor necrosis factor – alpha (TNFα), which is necessary for homeostatic synaptic scaling up of glutamate synapses on cultured hippocampal neurons^16^.

CRH^PVN^ neurons are a key node in the neural stress circuitry. Once active, these neurons initiate the production of CORT via the HPA axis, which is an essential component of an appropriate stress response. Recent observations show these neurons have functions beyond endocrine control^2,24,41,42^. Since these neurons have important roles in many different outputs, a mechanism of escaping persisting feedback may be an adaptive means of maintaining their influence over their various functions. It is interesting that the CORT solution blunted HPA output following FST despite sustained CRH^PVN^ neuron activity. There are many possible explanations for this. First, CORT feedback has been shown to reduce adrenal weight^20^, suggesting an inability to produce CORT upon stimulation. Second, CORT feedback can decrease expression of CRH mRNA in the PVN^9^. Therefore, even if CRH^PVN^ neurons were active, a lack of CRH release would prevent downstream HPA activation.

An adaptive increase in the strength of glutamate allows CRH^PVN^ neurons to maintain basal activity during sustained CORT feedback. This suggests that these neurons are not just responding, but continually adjusting to changes that are imposed by stimuli that threaten activity. It is possible that these adaptations may eventually result in an allostatic load on the system. If so, changes in this adaptive regime may provide a fruitful approach when targeting network disruptions in chronic disease or psychopathology. Indeed, observations that adjustments in synaptic strength is a target for mood correcting drugs such as lithium^43^ is consistent with this idea. It is also interesting to note that pharmacotherapy that corrects elevated CORT levels in diseases such as Cushing’s, does not always resolve the behavioral symptomology.

## Acknowledgements

We thank Ms. Cheryl Sank for technical assistance and Ms. Mio Tsutsui for microinjections and histology. We thank the Advanced Light and Optogenetics facility at the Cumming School of Medicine for access to photometry tools. We are grateful to members of the Bains lab for many helpful discussions on earlier versions of this manuscript.

## Author Contributions

N.P.R. and J.S.B. designed all experiments and prepared the manuscript. N.P.R. conducted the experiments, analyzed the data, prepared figure and wrote the manuscript. J.S.B. supervised the entire project, prepared figures and wrote the manuscript. T.F. and D.G.R. built the fiber photometry system. J.M. and D.G.R. designed the custom analysis software for fiber photometry data analysis. N.D. assisted with procuring plasma and conducting corticosterone ELISA assays. T.L.S. assisted with experiments involving patch-clamp electrophysiology.

## Declarations

The authors have no declarations to report.

## Methods

### Animals and CORT/DHF treatment

CRH-CretdTomato mice were used in this study. Animals were singly housed in a colony room with 12:12hr light-dark cycle. Mice were given ad libitum access to a 0.95% EtOH solution containing 25μg/ml CORT as their sole source of drinking water for 7 days. To prepare the solution, corticosterone (Sigma), was first dissolved in 95% ethanol and sonicated for 1-2min before dilution with tap water to 25μg/ml. For experiments involving 7,8 dihydroxyflavone (DHF), DHF was also dissolved in 95% EtOH and sonicated for 1-2min before being diluted in tap water. For electrophysiological experiments, mice were aged p30 – p40. Mice used for in vivo recording were older (p56 – p70), as they required time to recover from surgical implantation. Solutions were replaced every 2-3 days and the amount of solution consumed was recorded. All water bottles containing solution were wrapped with foil to prevent light exposure.

### Electrophysiology

Mice were anesthetized with isoflurane and decapitated. The brain was rapidly extracted and submerged in freezing slicing solution (0 °C, 95% O_2_, 5% CO_2_ saturated) containing 87 mM NaCl, 2.5 mM KCl, 0.5 mM CaCl2, 7 mM MgCl2, 25 mM NaHCO3, 25 mM D-glucose, 1.25 mM NaH2PO4 and 75 mM sucrose. Coronal slices were taken (250μm) using a vibratome (Leica). The PVN was hemisected and slices were incubated for 1 hour in artificial cerebral spinal fluid (aCSF) (30 °C, 95% O_2_, 5% CO_2_ saturated) containing 126 mM NaCl, 2.5 mM KCl, 26 mM NaHCO3, 2.5 mM CaCl2, 1.5 mM MgCl2, 1.25 mM NaH2PO4 and 10 mM glucose. Once transferred to a recording chamber superfused with aCSF (1 ml min−1, 30–32 °C, 95% O_2_, 5% CO_2_), slices were visualized using an Olympus BX51WI upright microscope fitted with infrared differential interference contrast optics. Pulled borosilicate glass pipettes (3–6 MΩ tip resistance) were filled with internal solution containing 108 mM potassium gluconate, 2 mM MgCl2, 8 mM sodium gluconate, 8 mM KCl, 1 mM K2-EGTA, 4 mM K2-ATP, 0.3 mM Na3-GTP and 10 mM HEPES. CRH^PVN^ neurons were identified by CRE-dependent expression of tdtomato fluorescent marker (previously characterized by Wamsteeker-Cusulin & Füzesi et al., 2013).

For voltage clamp recordings, mEPSCs were recorded for 3min at −63 mV in aCSF solution containing TTX (1μM, Alamone) and picrotoxin (100 μM, Sigma). mEPSCs were analyzed with Minianalysis, (Synaptosoft), and a detection threshold of 8.5pA was used based on 5x RMS noise (RMS = 1.7pA). Spontaneous inhibitory postsynaptic currents were recorded at 0mV in the absence of TTX or ion channel antagonists for 3min. Minianalysis, (Synaptosoft) was used to analyze sIPSCs, and a detection threshold of 35pA was used. The membrane capacitance and membrane resistance recorded shortly after gaining whole-cell access. For experiments requiring evoked EPSCs, AMPA currents were electrically evoked at −70mV in picrotoxin (100μM, Sigma) with a stimulating electrode located in the paraventricular aspect (Marty et al., 2011). PPR was attained using a paired stimulation (50ms apart at 0.1Hz intervals). Evoked currents were analyzed with Clampfit (Molecular Devices). For all voltage clamp recordings, access resistance was monitored, and cells were discarded if access resistance was >20MΩ or if access changed >15% during recording. For current clamp experiments, cells were held at −70mV, and incremental current steps of 10pA (from - 40pA to 120pA, at 0.5s) were delivered at 1Hz.

For experiments involving γ-DGG, mEPSCs were first recorded for 3min. Then, γ-DGG was infused into the bath (bath concentration was 500µM) for 5 min, and mEPSCs were recorded for another 3min. The first 150 events were normalized to the largest event before and after the antagonist was added to generate the cumulative distributions. A ratio of mean amplitude before and after addition of the antagonist was calculated for comparison.

### Synaptic scaling analysis

To assess for synaptic scaling, the first 250 mEPSCs recorded from 12 cells in each condition were used. This yielded a population of 3000 events from each condition, which were then compiled into a cumulative distribution. We then used a method characterized by Kim et al., 2012 to assess for synaptic scaling. The amplitude distributions from CORT treated animals were divided by several factors to achieve the highest P-value when compared against the distribution from naïve animals using the Kolmogorov-Smirnov (KS) test. As per the method of Kim et al., 2012, as the distributions from CORT treated animals were scaled back, mEPSCs on the lower end of the distribution that fell below the event detection threshold (8.5pA) were discarded, as this can skew the K-S test. For the rank-order plot, events within the 5th – 95th percentile were used to generate the graph and linear regression.

### Corticosterone quantification and forced swim stress

For CORT quantification experiments, mice were habituated to handling 3 days prior to the experiment, and an initial tail vein cut was performed at the distal portion of the tail at the end of habituation on the 3rd day. Plasma was extracted by reopening the wound with a surgical blade. A microvette tube (CB 300 Z, Sarstedt AG & Co) was used to collect the plasma, before being centrifuged (8,000RPM for 20min, at 4°C). The plasma was separated from the hematocrit and was stored in an Eppendorf tube at −20°C until further analysis. DetectX Corticosterone Immunoassay Kit was used to detect plasma-[CORT] (Arbor Assay). Samples were assessed in triplicate and averaged. For FST experiments, animals were placed in a 1L beaker filled with 36°C water for 15min, and the water was changed between animals. Tail vein Plasma was collected 2hrs prior for baseline comparison and immediately following the FST.

### Viral injection and fiber photometry

Crh-IRES-Cre;Ai14 (p35-42) were unilaterally injected with AAV9-CAG-Flex-GCaMP6s (Penn Vector Core). Mice were placed in a stereotactic apparatus and anesthetized with 4% isoflurane. A glass capillary containing the virus was lowered into the brain (coordinates from bregma: bilateral ± 0.2mm, anterior-posterior 0.0mm, dorsal-ventral - 4.6mm). The virus was pressure injected using the Nanoject II (Drummond Scientific). Animals received 3 injections of 69nL to each lateral hemisphere. To prevent backflow of the virus, we allowed 2min between injection for the first 2 injections, and 10min after the last injection before retracting the injection capillary. Mice were given 2 weeks recovery.

After 2-week recovery, the mice had a mono-fiber cannula (Doric MFC_400/430-0.48_5mm) implanted ~100µm dorsal to PVN, and were given at least 7 days to recover. Animals were then habituated to handling and fiber attachment (30min/day) for 4 days prior to beginning experiments. GCaMP6s signal was collected using a photometry rig equipped with an LED controller (Thor Labs) that generated a 470nm wavelength (211Hz) for GCaMP6s signal, and a 405nm wavelength (530Hz) that was collected as a reference signal to correct for motion artifact. For excitation of GCaMP6s, the wavelengths first passed through a filterset (Doric), before going through a 400µm core-diameter fiber (Doric Lenses), which was then connected to the cannula on the animal. The power of each wavelength at the tip of the fiber was calibrated to 30µW before each recording session. The signals generated by GCaMP6s was collected by the fiber, and passed through the filterset and photo-receiver (Newport). The signal entered an amplifier (Tucker Davis Technologies) for demodulation, and was digitized at 1.0173 kHz. The signal was recorded using Synapse Software.

Photometry data were analyzed using custom MATLAB scripts. Both traces were fit to a 2^nd^ order polynomial to correct for bleaching. After fitting, the reference signal was subtracted from the 470nm signal. We generated all-points histograms from these traces by decimating the trace by a factor of 100. Then, 60s of baseline (611 data points) from each animal were plotted on a histogram to yield a distribution around the mean and compared with K-S statistics. For peak detection, a modified script from Muir et al., 2018 was used. Both the 470nm and 405nm signals were plotted against each other and a linear regression was generated. The reference (405nm) wavelength was scaled by the slope of the linear regression, and then subtracted from the GCaMP6s (470nm) signal. Then, the resultant trace was high-pass filtered (0.1Hz cut off). Events larger than 2x MAD were counted as peaks.

### Data analysis and statistics

Statistical analysis was performed using GraphPad Prism. Data was presented as mean ± standard error of the mean.

---

**Fig. S1.**
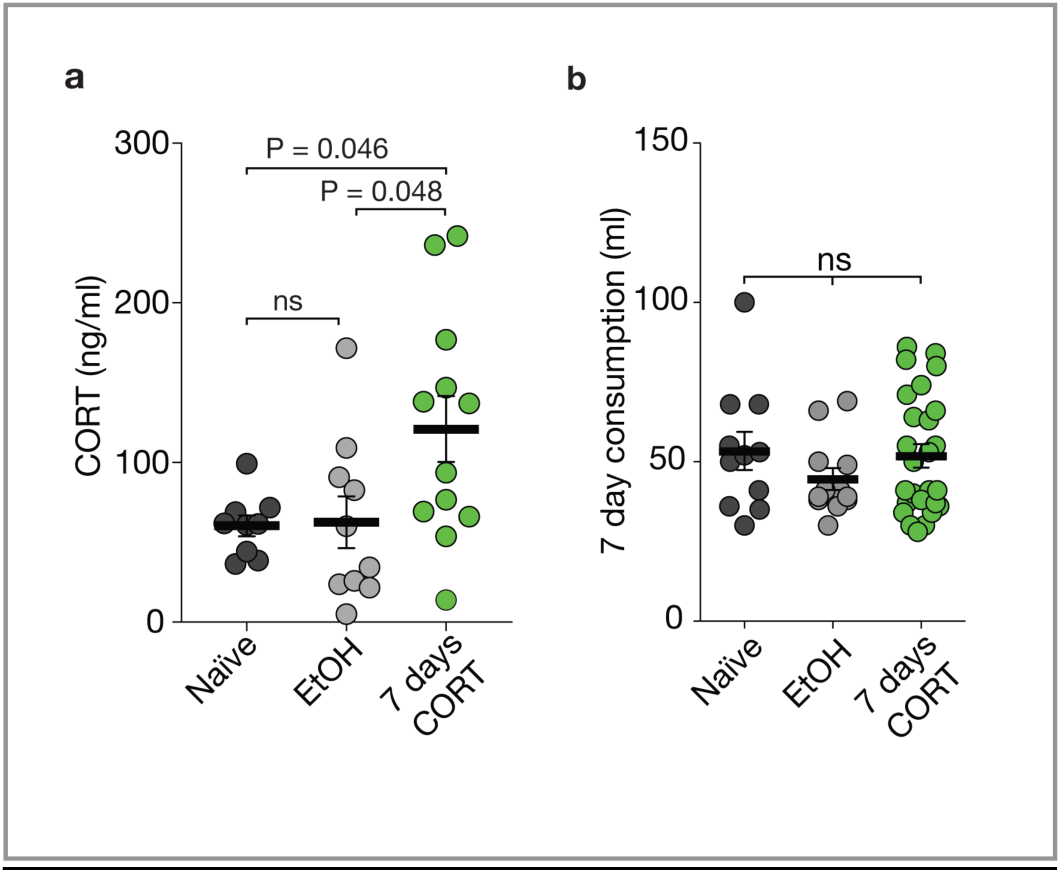
a) Mice drinking exogenous CORT had elevated circulating CORT levels during the dark phase when compared to naïve and EtOH (naïve, 60.43 ± 6.49ng/ml, N = 9 vs 7 days CORT, 121.0 ± 20.72ng/ml, N = 12, vs EtOH, 62.58 ± 16.31ng/ml, N = 10, one-way ANOVA, Tukey correction, F(2, 28) = 4.34, P = 0.023). b) There was no difference in the amount of solution consumed over 7 days (naïve, 53.45 ± 6.00ml, N = 11, vs EtOH, 44.67 ± 3.44ml, N = 12, vs, CORT, 51.92 ± 3.68ml, N = 26, Kruskal-Wallis test, Kruskal-Wallis statistic = 1.021, P = 0.60).

**Fig. S2.**
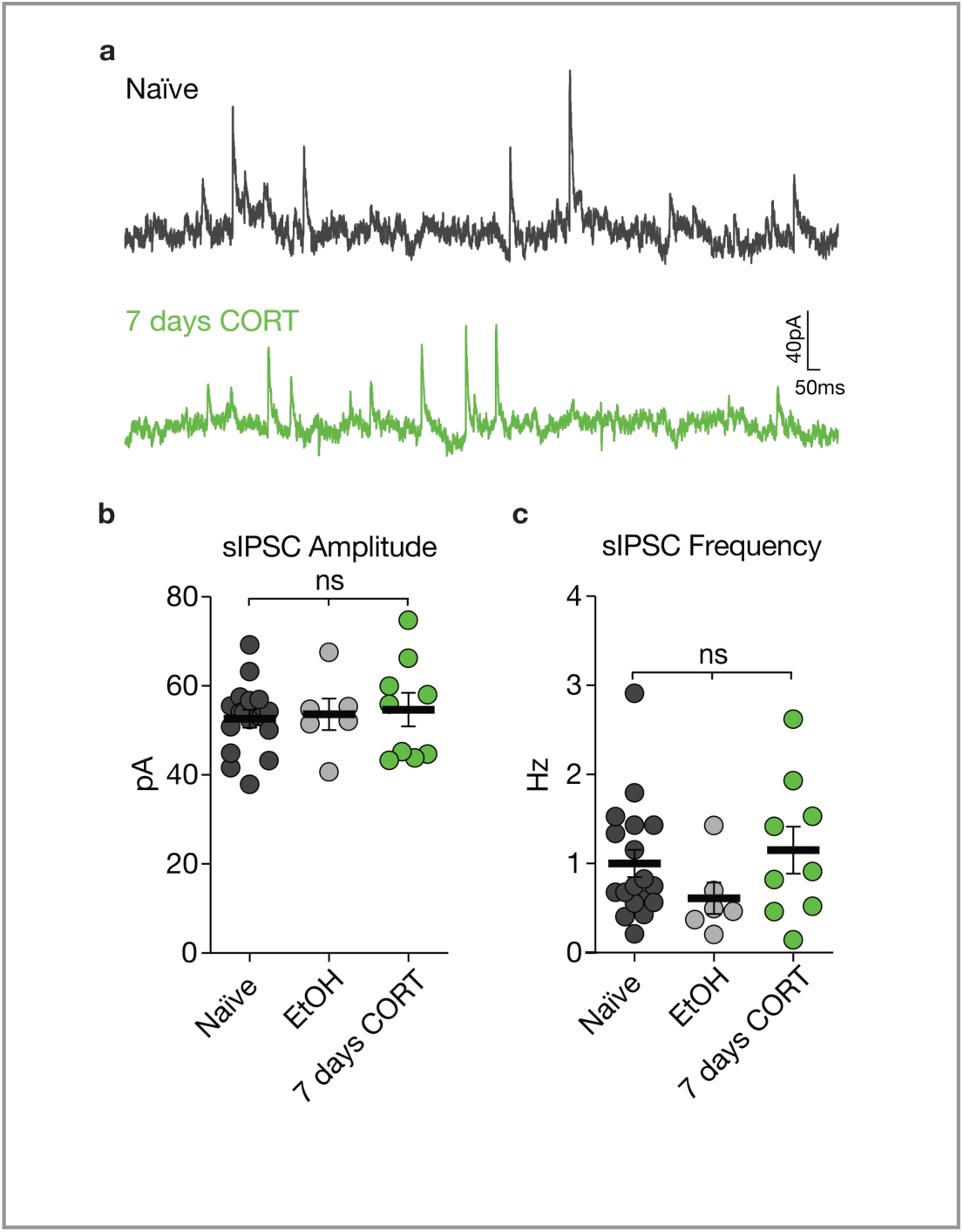
a) Representative traces recorded at 0mV holding potential. b) There was no change in sIPSC amplitude (naïve, 52.6 ± 1.99pA, n = 16, vs, EtOH, 53.7 ± 3.53pA, n = 6, vs 7 days CORT, 54.2 ± 3.68pA, n = 9, one-way ANOVA, Tukey correction, F (2, 28) = 0.0910, P = 0.91). c) There was no change in sIPSC frequency (naïve, 1.04 ± 0.171Hz, n = 16, vs EtOH, 0.611 ± 0.176Hz, n = 6, vs 7 days CORT, 1.47 ± 0.401Hz, n = 9, Kruskal-Wallis test, Dunn’s correction, K-W statistic = 3.57, P = 0.17).

**Fig. S3.**
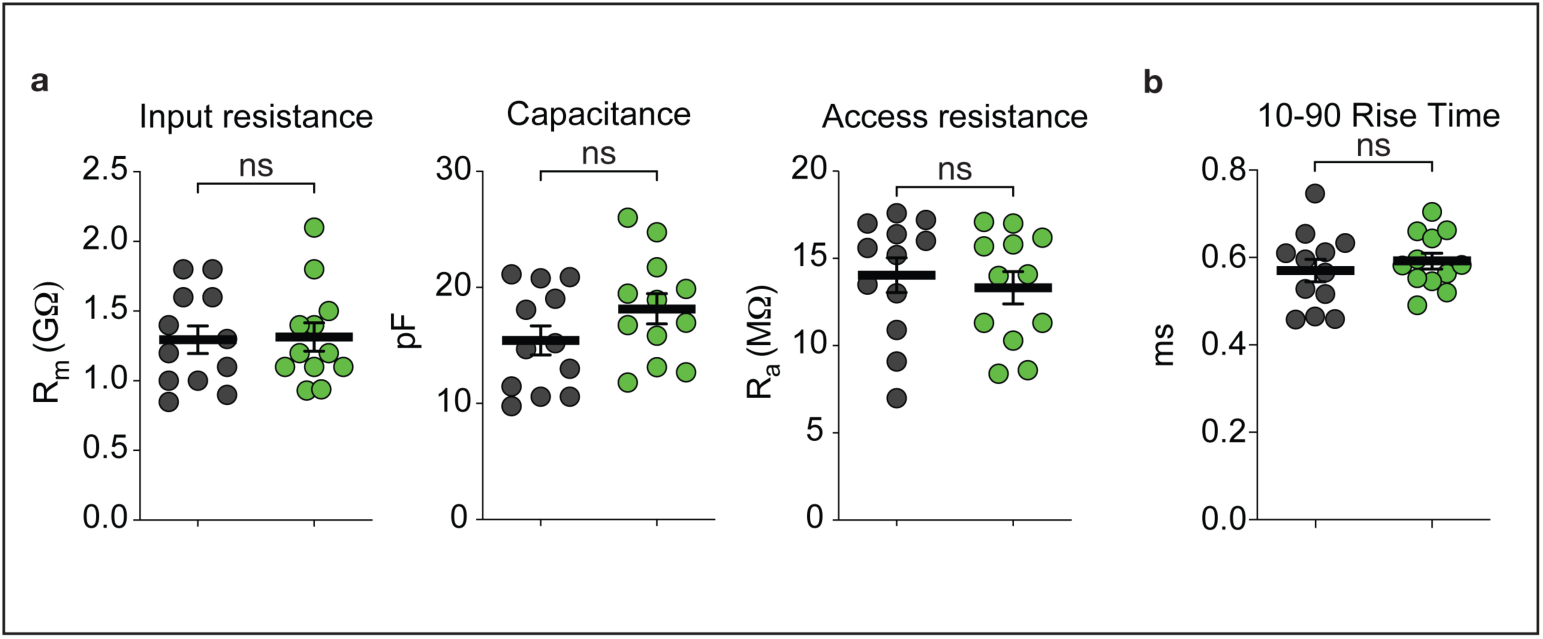
CORT did not change input resistance, capacitance, or access resistance. Input resistance, naïve, 1.30 ± 0.099GΩ vs 7 days CORT 1.31 ± 0.10GΩ, n = 12, two-tail student’s t-test, t(22) = 0.13, P = 0.89. Access resistance, naïve, 14.04 ± 0.99MΩ vs 7 days CORT, 13.32 ± 0.92MΩ, n = 12, two-tail student’s t-test, t(22) = 0.53, P = 0.60. Capacitance, naïve, 15.44±1.27pF vs 7 days CORT, 18.17 ± 1.13pF, n = 12, student’s t-test, t(22) = 1.49, P = 0.15. b) 10-90 rise time was not changed by CORT (naïve, 0.57 ± 0.026ms vs 7 days CORT, 0.59 ± 0.018ms, n = 12, two-tail student’s t-test, t(22) = 0.68, P = 0.50).

**Fig. S4.**
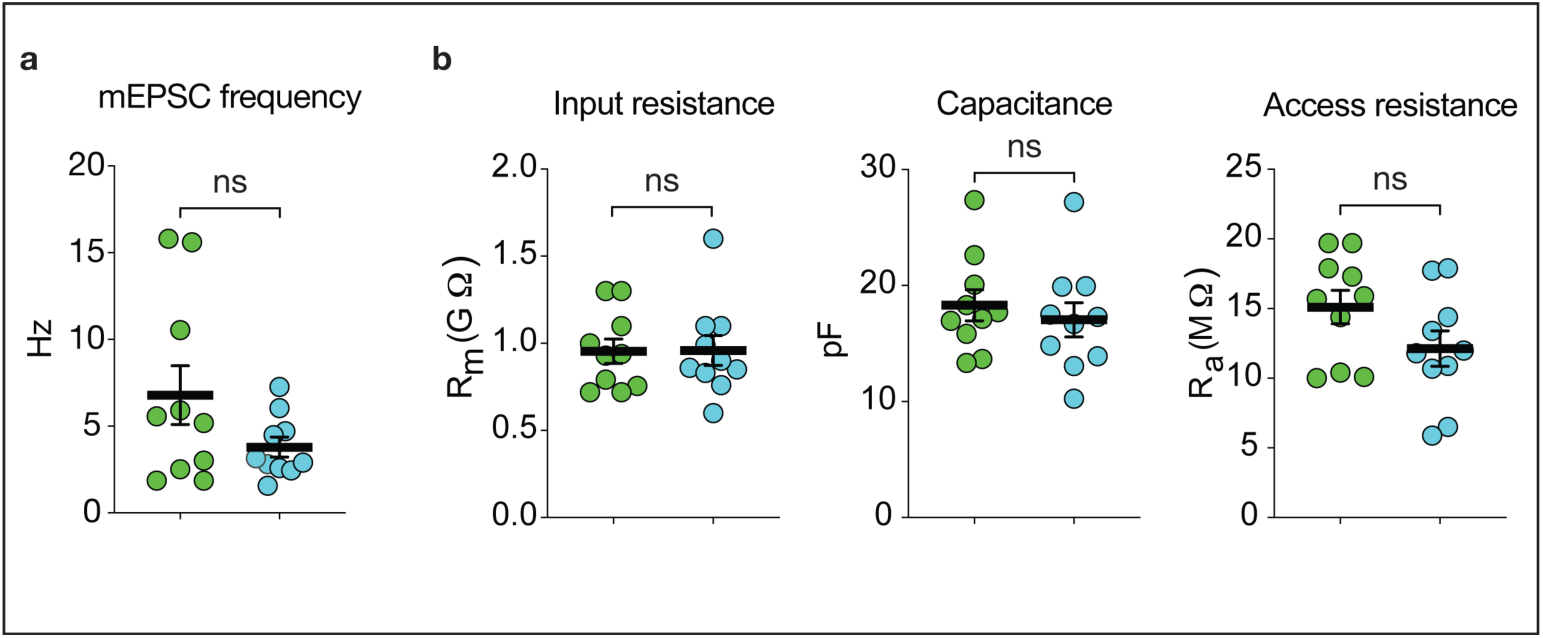
a) There was no difference in mEPSC frequency (CORT, 6.79 ± 1.70Hz vs CORT + DHF, 3.80 ± 0.566Hz, n = 10, two-tail student’s t-test, t(18) = 1.67, P = 0.11). b) There was no difference in the input resistance, capacitance, or the access resistance. Input resistance, CORT, 0.956 ± 0.0698GΩ vs CORT + DHF 0.959 ± 0.0858GΩ, n = 10, two-tail student’s t-test, t(=18) = 0.0307, P = 0.98. Capacitance, CORT, 18.3 ± 1.33pF vs CORT + DHF, 17.1 ± 1.48pF, n = 10, student’s t-test, t(18) = 0.626, P = 0.15. Access resistance, CORT, 15.1 ± 0.1.20MΩ vs CORT + DHF, 12.1 ± 1.27MΩ, n = 10, two-tail student’s t-test, t(18) = 1.71, P = 0.10. b)

## Notes

### Competing Interest Statement

The authors have declared no competing interest.

